# A warming Southern Ocean may compromise Antarctic blue whale foetus growth

**DOI:** 10.1101/2020.06.25.170944

**Authors:** Carl Smith

## Abstract

After declining in abundance due to commercial whaling during the 20^th^ Century, populations of the Antarctic blue whale (*Balaenoptera musculus intermedia*) have failed to recover to pre-exploitation levels. Using historical whaling data from 1926-1954, in combination with temperature data for the Southern Ocean, a gamma GLMM with temporal dependency was fitted to 20,144 records of *B. m. intermedia* foetus size using Bayesian inference. There was a negative relationship between antecedent winter sea surface temperature (SST) in the Southern Ocean on foetus size. This relationship is proposed as being mediated by a positive effect of the extent of winter sea ice on Antarctic krill (*Euphasia superba*) abundance on which *B. m. intermedia* feed. There was also a positive density-dependent effect of a ‘krill surplus’ at low whale population sizes. However, the positive effect of a ‘krill surplus’ at low *B. m. intermedia* population size on foetus growth was reversed at elevated winter SST due to a negative impact on *E. superba* recruitment. Projected increases in temperature in the Southern Ocean are predicted to compromise the growth rates of *B. m. intermedia* foetuses, with implications for the capacity of the subspecies to recover from overexploitation.

## Introduction

The large-scale harvest of baleen whales (Balaenopteridae) in the Southern Ocean, which began at the start of the 20^th^ Century, saw rapid declines in the abundance of blue (*Balaenoptera musculus*), humpback (*Megaptera novaeangliae*), fin (*B. physalus*) and sei (*B. borealis*) whales (Baker & Clapham, 2004). After several decades of protection, populations of these whales have shown varying degrees of recovery (Best, 1993; Branch, Matsuoka, & Miyashita, 2004; Clapham, Young, & Brownell, 1999). *M. novaeangliae* have shown strong recovery, *B. physalus* populations have broadly shown increases in numbers, while the status of *B. borealis* populations is poorly known (Thomas, Reeves, & Brownell, 2016). *B. musculus* populations show wide variation in status and recovery; some populations have disappeared entirely or remain at extremely low levels (Thomas et al., 2016), while at least one population from the eastern North Pacific appears to have recovered to pre-exploitation levels (Monnahan, Branch, Punt, 2015). The Antarctic blue whale (*B. m. intermedia*) is classified as Critically Endangered (Reilly et al., 2008). Prior to the onset of commercial whaling the abundance of *B. m. intermedia* was estimated at approximately 239,000 individuals. Abundance declined to as few as 370 individuals, despite a moratorium on the hunting of this species in 1965 (Branch et al., 2004). The most recent conservative population estimate for this subspecies was 2,280 individuals, barely 1% of pre-exploitation levels (Branch, 2007).

The underlying reasons for the slow recovery of *B. m. intermedia* have not been identified but may, in part, arise from illegal hunting by the Soviet Union into the 1970s, as well as from on-going mortalities associated with boat strikes and entanglement in fishing gear (Clapham et al., 1999; Yablakov, 1994). An additional explanation for their slow recovery could relate to a decline in food availability, particularly of Antarctic krill (*Euphasia superba*) on which *B. m. intermedia* feed almost exclusively (Bannister, 2008). In the southern hemisphere, *B. m. intermedia* migrate to Antarctic waters to feed during the austral summer, before migrating back to lower latitudes for the remainder of the year, where feeding is typically more limited, though there are departures from this general pattern (Hucke-Gaete et al., 2018). The period spent feeding is critical (Mackintosh, 1965; Hucke-Gaete et al., 2018), particularly as females arrive in the Southern Ocean pregnant, with foetal growth contingent on the accumulation of energy reserves by the mother on the feeding grounds (Christiansen, Víkingsson, Rasmussen, & Lusseau, 2014).

*E. superba* are an important species in the Southern Ocean (Fraser & Hofmann, 2003), feeding on blooms of phytoplankton at the edge of the Antarctic sea ice with their abundance closely linked to the extent of ice (Braithwaite, Meeuwig, Letessier, Jenner, & Brierley, 2015; Brierly et al., 2002). Declines in the extent and duration of Antarctic winter sea ice since the 1970s have resulted in a decline in the abundance of *E. superba* (Atkinson, Siegel, Pakhomov, & Rothery, 2004; Nicol, 2006), with potential consequences for the predators that rely on them, including baleen whales (Ainley et al., 2006, 2007; Alonzo & Mangal, 2001; Reid & Croxall, 2001). Braithwaite et al. (2015) linked the extent of sea ice with the quantity of oil extracted from the carcasses of humpback whales caught during commercial whaling between 1947-1963. They showed that the extent of winter sea ice correlated positively with oil yields, suggesting that *M. novaeangliae* body condition was directly linked to *E. superba* abundance, mediated by the extent of sea ice.

Balaenopterid whales display the highest foetal growth rates of all mammals (Frazer & Huggett, 1973), demanding substantial allocation of maternal somatic resources for embryonic and foetal growth (Lockyer, 1986). Rapid foetal growth in marine mammals permits the production of large neonates with high survival rates (Boltnev, York, & Antonelis, 1998; McMahon, Burton, & Bester, 2003) and in the case of balaenopterid whales is synchronised with their annual feeding migration from low to high latitudes (Chivers, 2008). Energy supplied to the developing foetus comes from reserves accumulated during the intense feeding phase of their annual migratory cycle. If female condition is compromised, the outcome is a decline in fertility (Williams et al., 2013) and reduced investment in embryonic and foetal growth (Christiansen et al., 2014), with potential negative consequences for calf survival (Lockyer, 2007).

Declines in balaenopterid population size due to whaling can release resources (a ‘krill surplus’) to survivors and thereby facilitate recovery through density-dependent effects (Ainley et al., 2007; Best, 1993; Fraser, Trivelpiece, Ainley, & Trivelpiece, 1992; Laws, 1977). Density-dependent population growth has been demonstrated in *B. physalus* (Williams et al., 2013). However, while there is some evidence for the modest recovery of *B. m. intermedia* (Branch, 2008), the continued low abundance of this species means that it remains critically endangered.

Here I indirectly test whether foetal growth of *B. m. intermedia* is determined by the availability of *E. superba* by examining the effect of temperature on *B. m. intermedia* foetal body size. A prediction is that a reduction in the extent of sea ice in response to elevated temperatures in the Southern Ocean will suppress *E. superba* productivity thereby compromising *B. m. intermedia* foetus development with negative consequences for their subsequent post-natal survival to maturity. An additional prediction is that a decline in the abundance of *B. m. intermedia* will result in a ‘krill surplus’, detectable as increased foetus growth rate. A further prediction is that these processes are linked, with temperatures in the Southern Ocean interacting with the effect of adult *B. m. intermedia* abundance. Thus, a combination of low winter temperatures combined with low *B. m. intermedia* abundance are predicted to enhance foetal growth rates while the converse, elevated winter temperatures and high adult abundances, would tend to supress foetal growth as a consequence of reduced krill availability. I test these scenarios using historical whaling records and temperature data to model *B. m. intermedia* foetus size in response to sea surface temperature (SST) and *B. m. intermedia* abundance.

## Methods

### Data

Blue whale data came from International Whaling Statistics (IWS) database and comprised measurements of female body length and foetus length, whether the foetus was single or a twin, along with the date of capture (www.iwc.int). The complete dataset included 20,144 complete records spanning 30 years from 1925-1954 with data grouped by Antarctic whaling season, spanning the austral summer (October - March). Data for the whaling seasons 1940 - 1946 were missing due to limited hunting during WWII and its immediate aftermath. SST anomaly data for the Southern Ocean came from the HadISST1 data set, which comprises a globally complete SST dataset from 1871 onwards at a spatial resolution of 1° latitude by 1° longitude, compiled by the UK Met Office Hadley Centre (www.metoffice.gov.uk/hadobs/hadisst) (details in Rayner et al., 2003). A nominal day of conception for *B. m. intermedia* of 1^st^ August was used to estimate the day of gestation on which a female was killed and its foetus collected. This date was based on an analysis by Roston, Lickorish, & Buchholtz (2013), which used previous published studies combined with IWS data and assuming a gestation period of 350 days. While the evidence indicates this date represents a peak in conception frequency, there is a potentially wide temporal window in the timing of conception in this species. While time-series data for *E. superba* abundance in the Southern Ocean are available (Atkinson et al., 2017) and could potentially be used to link food abundance with foetus size, for the period of this study these data are incomplete and were not collected using consistent methodologies.

Density-dependent effects on foetus size were modelled using the total cumulative catch of *B. m. intermedia* in the whaling season preceding the one in which a foetus was collected. *B. m. intermedia* catch data came from Branch et al. (2008) and included a correction for Soviet misreporting of catches. Catch data, rather than estimated population size, was used as an inverse index of abundance since catch data were relatively reliably recorded, while estimates of population size necessarily include a margin of error (Christensen, 2006).

### Data analysis

Before applying statistical models, a data exploration was undertaken following the protocol described in Ieno & Zuur (2015). The data were examined for outliers in the response and explanatory variables, homogeneity and zero inflation in the response variable and the nature of relationships between the response and explanatory variables. Collinearity between explanatory variables was examined using co-plots and by estimating variance inflation (VIF). Model covariates were not collinear and VIF of covariates in the model were all <2. Model assumptions were verified by plotting model residuals against fitted values, each covariate in the model and covariates not included in the model. Model residuals were additionally assessed for evidence of non-linearity (Zuur & Ieno, 2016). Twin foetuses, of which there were 312 in the complete dataset, were consistently smaller that singletons and were dropped from the analysis. If twin foetuses were retained in the model the qualitative outcomes were identical.

Data were modelled using R (version 3.6.3; R Development Core Team, 2020) with models fitted in a Bayesian framework using Integrated Nested Laplace Approximation (R-INLA; Rue et al., 2017). The advantage of using Bayesian inference was that it provides probability distributions for model parameters of interest, so that probability statements about the magnitude of model parameters can be made with confidence. This approach avoids reliance on hypothesis testing and P-values, which are increasingly recognised as unreliable statistical tools for any but the simplest models (Burnham & Anderson, 2014; Nuzzo, 2014; Wasserstein & Lazar, 2016).

To accommodate temporal dependency, foetus size was modelled with a gamma GLMM with a second order random walk trend fitted for day of gestation. The model was formulated as:

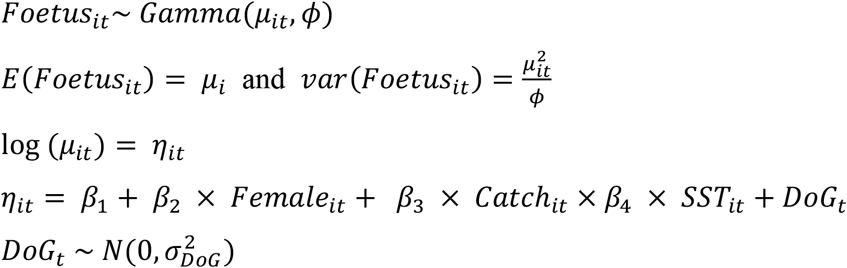

Where *Foetus*_*it*_ is the mean size (*μ*) of foetus *i* on day of gestation (*DoG*) *t*. The parameter *ϕ* is a dispersion parameter (Zuur, Ieno & Saveliev, 2017). The model contained linear effects for female length (*Female*_*it*_), cumulative *B. m. intermedia* catch in the whaling season preceding the one in which the foetus was collected (*Catch*_*it*_), and mean SST of the Southern Ocean for the preceding austral winter (April-September) (*SST*_*it*_). Non-informative priors were put on δ and τ, with Jeffreys prior (*a*_1_ = *b*_1_ = 0) on δ and a Gelman (2006) recommended prior (*a*_2_ = - 0.5, *b*_2_ = 0) on τ.

Because the window of conceptions for *B. m. intermedia* is potentially wide (Roston et al., 2013), any consistent environmental effects on the time of conception could undermine conclusions relating to effects of environmental temperature and foetus size. To accommodate this possibility, a model was also fitted to the difference between observed foetus size and fitted estimates of foetus size as a function of mean SST for the preceding austral winter using a Gaussian GLM, with the prediction that any relationship between foetus size and SST (either positive or negative) would indicate an association between conception date and antecedent winter SST.

## Results

There was a statistically important negative interaction between cumulative *B. m. intermedia* catch and SST anomaly on *B. m. intermedia* foetus size (Table 1; Fig. 1). Female *B. m. intermedia* body size was positively associated with foetus size, after controlling for variance in day of gestation, SST anomaly and *B. m. intermedia* catch (Table 1). The smoother for the random effect of day of gestation showed a strong positive temporal trend (Fig. 2).

**Table 1.**
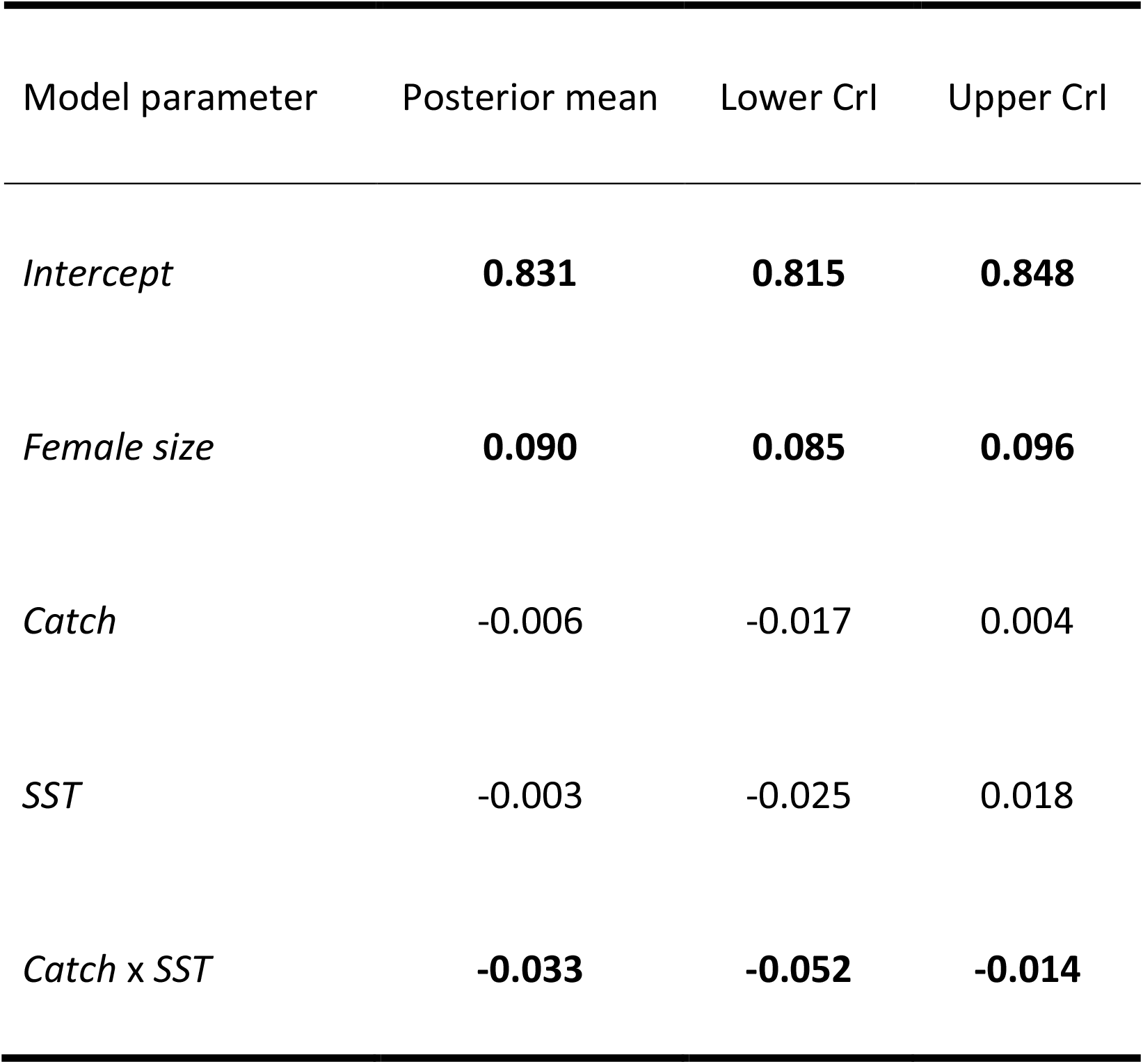
Posterior mean estimates of *B. m. intermedia* foetus size modelled using a random walk gamma GLMM fitted using INLA with non-informative priors. CrI is the 95% Bayesian credible interval. Credible intervals that do not contain zero in bold to indicate statistical importance. *Female size* is the length of female *B. m. intermedia* from which a foetus was taken. *SST* is the mean sea surface temperature anomaly of the Southern Ocean for the preceding austral winter (April-September). *Catch* is the cumulative catch of *B. m. intermedia* in the preceding whaling season.

| Model parameter | Posterior mean | Lower CrI | Upper CrI |
| --- | --- | --- | --- |
| *Intercept* | **0.831** | **0.815** | **0.848** |
| *Female size* | **0.090** | **0.085** | **0.096** |
| *Catch* | -0.006 | -0.017 | 0.004 |
| *SST* | -0.003 | -0.025 | 0.018 |
| *Catch x SST* | **-0.033** | **-0.052** | **-0.014** |

**Figure 1.**
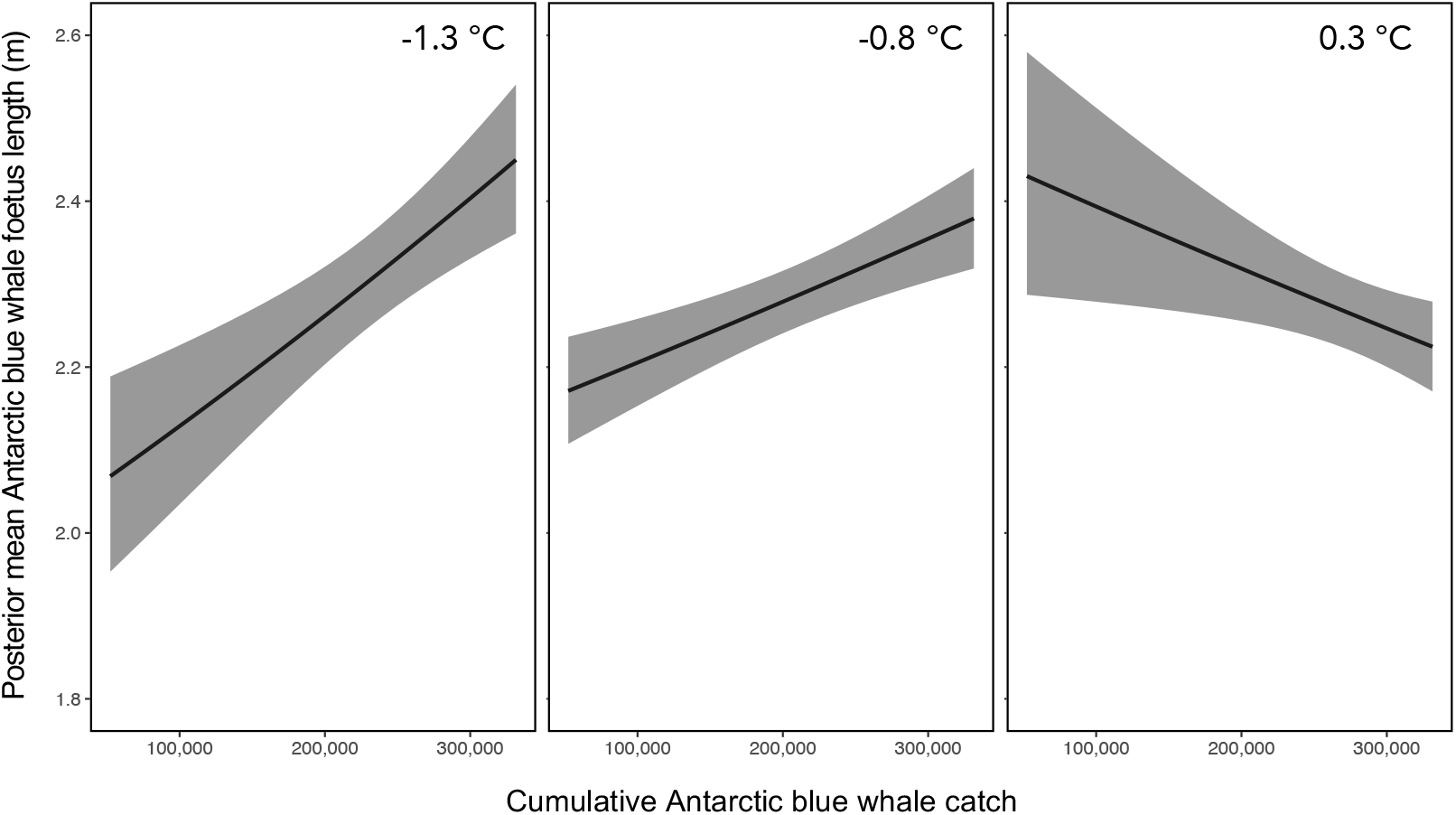
Posterior mean (solid line) fitted Antarctic blue whale (*Balaenoptera musculus intermedia*) foetus length (m), with 95% credible intervals (shaded area), as a function of the interaction between cumulative *B. m. intermedia* catch and antecedent winter sea surface temperature (SST) anomaly in the Southern Ocean, showing a reversal of the positive relationship between foetus length and catch at low SST (−1.3 °C, -0.8 °C), to a negative relationship at elevated SST (0.3 °C).

**Figure 2.**
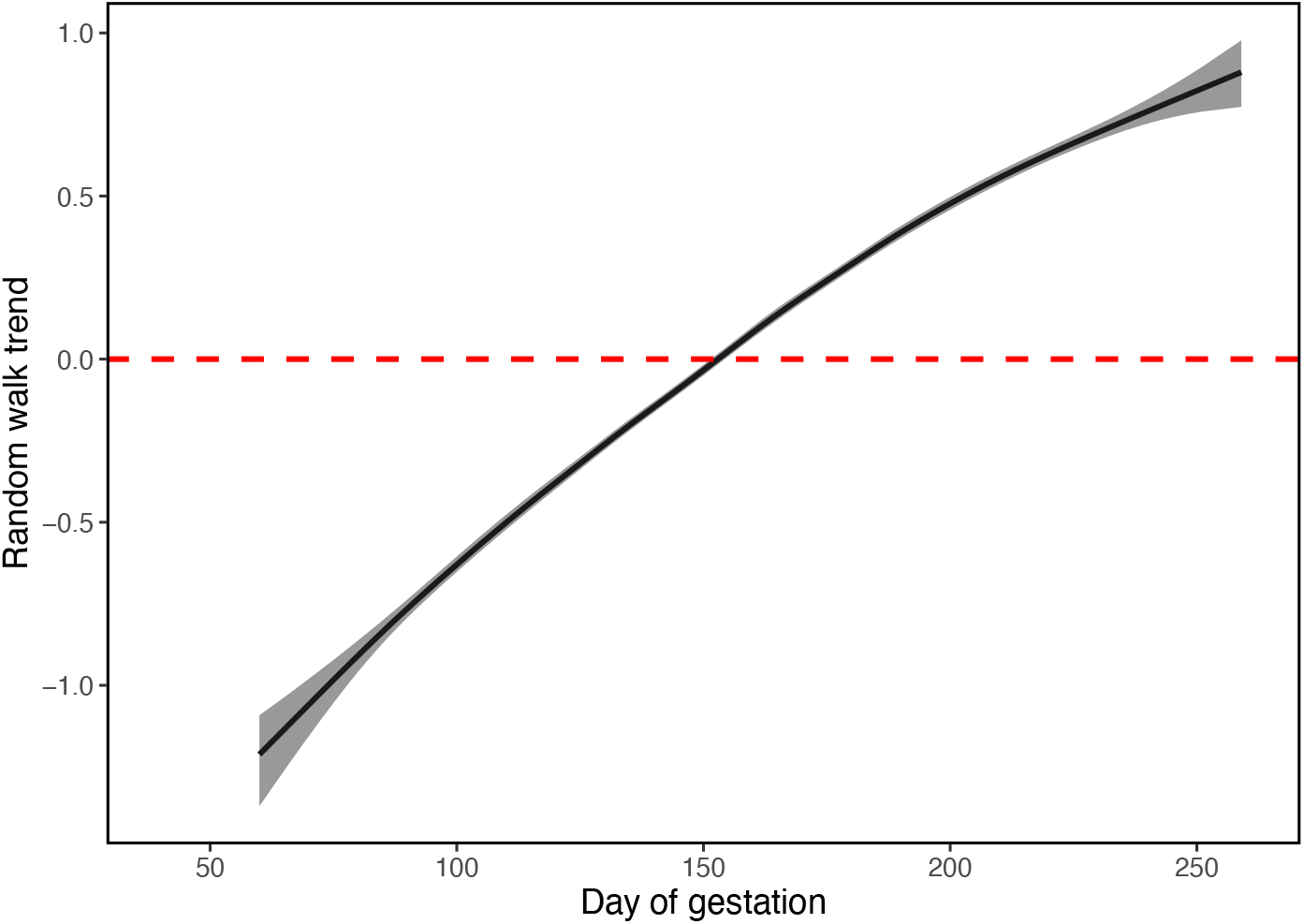
Posterior mean (solid line) with 95% credible intervals (shaded area) of the random effects of the random walk gamma GLMM for Antarctic blue whale (*Balaenoptera musculus intermedia*) foetus lengths against day of gestation assuming a nominal day of conception of 1^st^ August.

Model estimates of foetus size over time showed a comparable trend to observed data (Fig. 3), though with lower variance. There was no association between observed foetus size and fitted estimates of foetus size as a function of mean SST (Fig. 4) indicating that, while there was substantial variability around the nominal day of conception of 1^st^ August, there was no association between conception date and SST (Table 2). The body size of pregnant *B. m. intermedia* over the period for which data are presented showed no trend, either upward or downward (Fig. 5).

**Table 2.** Posterior mean estimates of the difference between observed foetus size and model estimates of foetus size as a function of antecedent winter sea surface temperature (*SST*) anomaly in the Southern Ocean modelled using a Gaussian GLM fitted using INLA with non-informative priors. CrI is the 95% Bayesian credible interval. If credible intervals do not contain zero it indicates statistical importance.

| Model parameter | Posterior mean | Lower CrI | Upper CrI |
| --- | --- | --- | --- |
| *Intercept* | 0.002 | -0.020 | 0.024 |
| *SST* | 0.006 | -0.037 | 0.049 |

**Figure 3.**
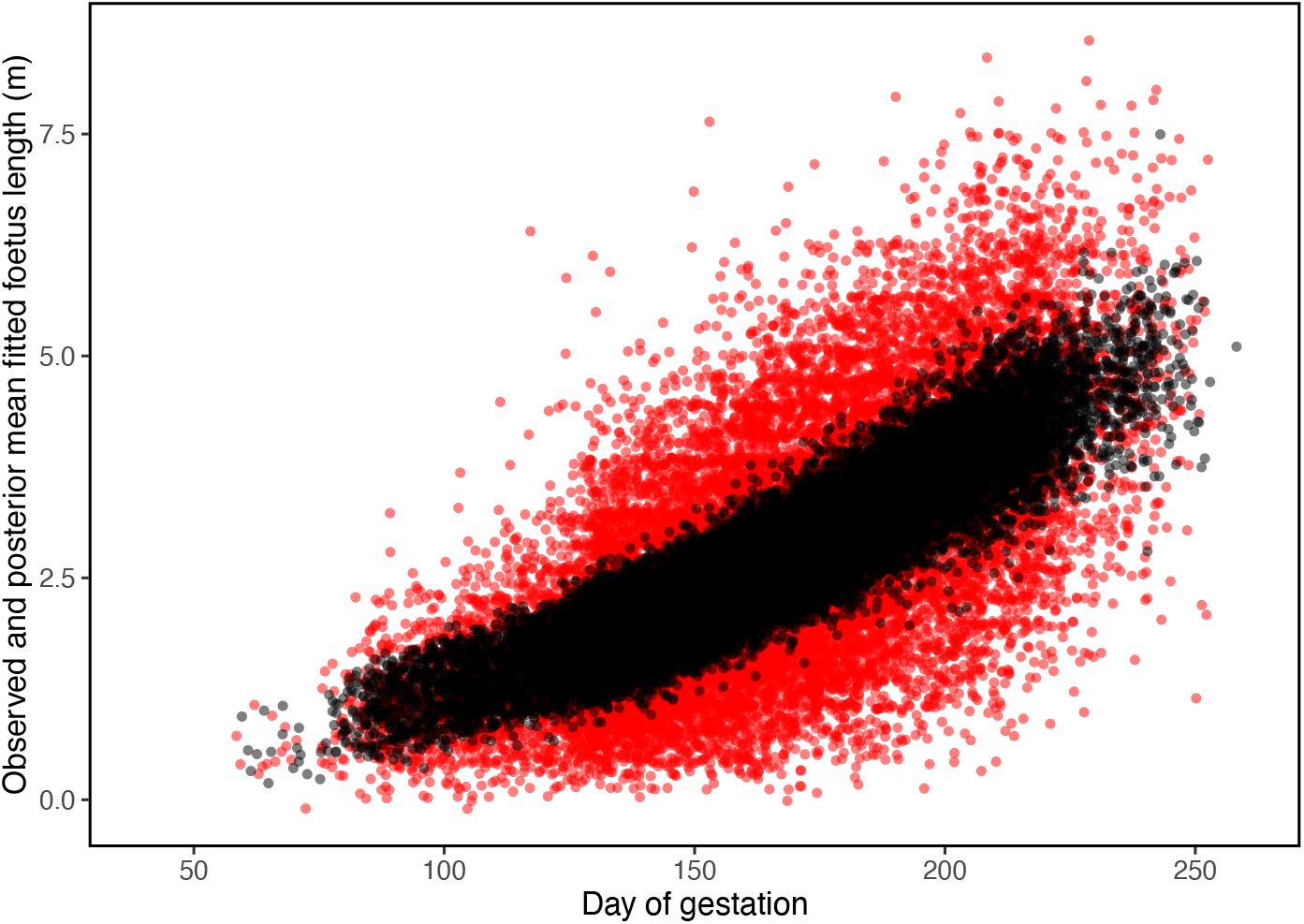
Plot of observed (red dots) and model fitted values (black dots) of Antarctic blue whale (*Balaenoptera musculus intermedia*) foetus lengths against day of year modelled with a random walk gamma GLMM assuming a nominal day of conception of 1^st^ August.

**Figure 4.**
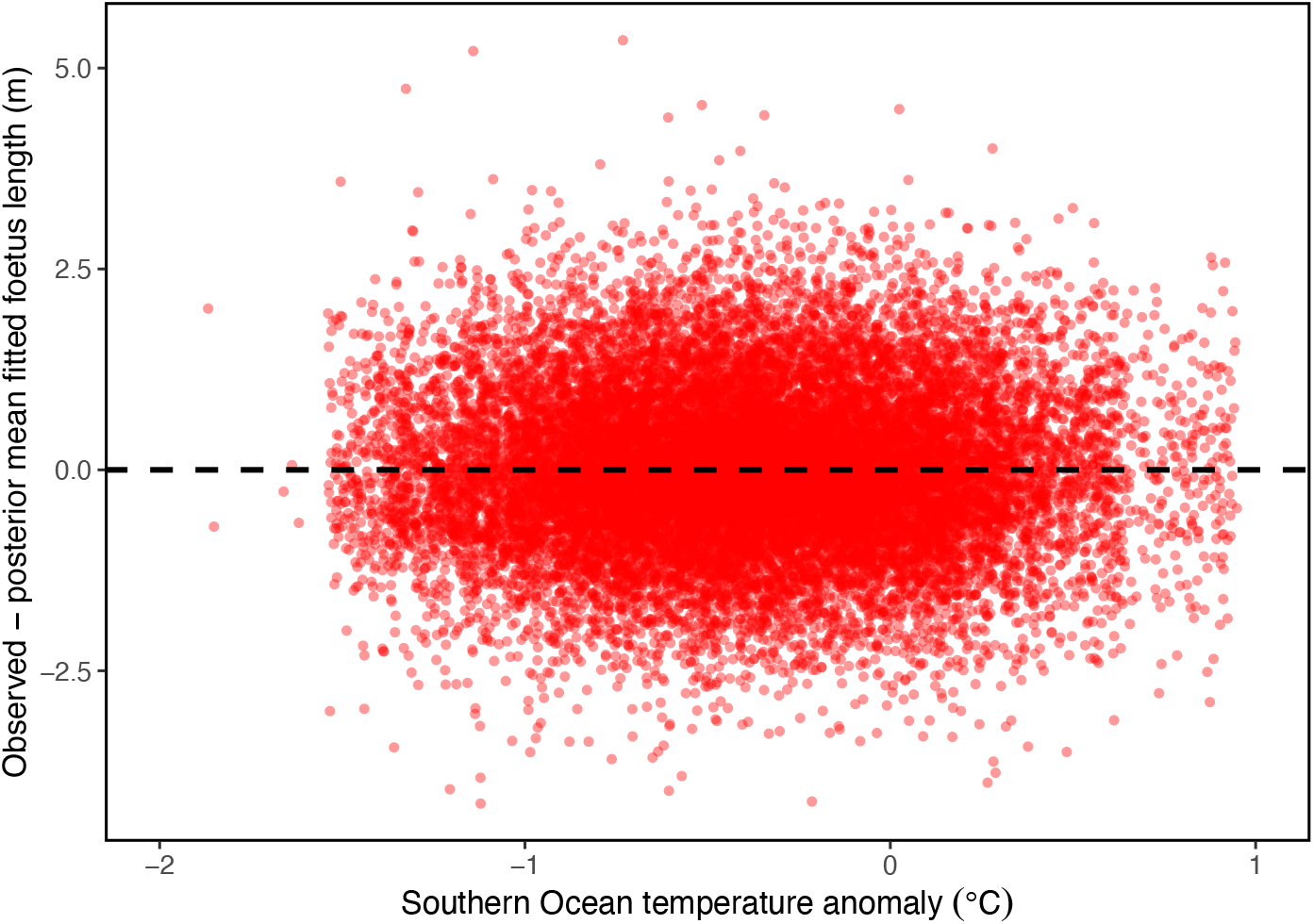
Difference between posterior fitted estimates of Antarctic blue whale (*Balaenoptera musculus intermedia*) foetus length (m) and observed foetus length as a function of average Southern Ocean temperature anomaly (°C) during the preceding austral winter (April – September). Horizontal black dashed line indicates unity.

**Figure 5.**
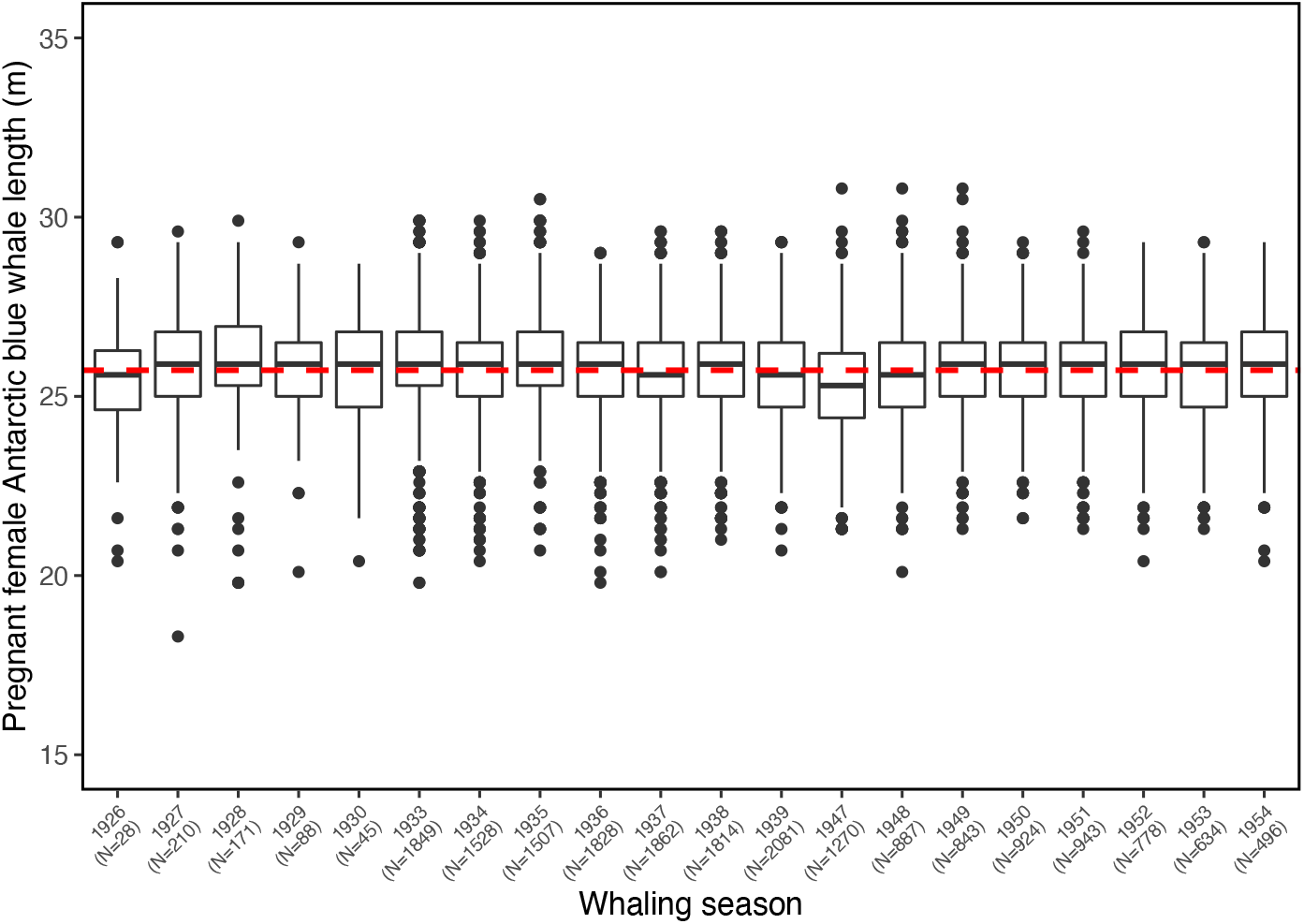
Boxplots of pregnant female Antarctic blue whale (*Balaenoptera musculus intermedia*) lengths (m) caught in the 1926 – 1954 whaling seasons. Horizontal red dashed line indicates mean female length. N is the number of females measured in each whaling season.

## Discussion

A statistically important negative interaction between SST anomaly in the Southern Ocean and abundance of adult *B. m. intermedia* on foetus growth was detected (Fig. 1, Table 1). Elevated winter temperatures in the Southern Ocean are associated with weak *E. superba* recruitment through limitation of the extent and duration of winter sea ice (Atkinson et al., 2004; Braithwaite et al., 2015; Brierly et al., 2002). A low abundance of *E. superba*, the main component of the diet of *B. m. intermedia* (Bannister, 2008), is predicted to have a negative effect on foetus growth. At the same time, the cumulative catch of *B. m. intermedia* through whaling was predicted to have a positive effect on foetus growth, through the greater abundance of *E. superba* as the population declined under hunting pressure (Fraser et al., 1992; Laws, 1977) (Fig. 1). The interaction between SST and *B. m. intermedia* catch on foetus size demonstrates that a positive effect of reduced blue whale density can be reversed at high SST (Fig. 1). An implication of this result is that while low Antarctic blue whale population size is predicted to result in increased foetal size-at-age through positive density dependence, increased temperatures in the Southern Ocean may offset any positive effects of reduced adult density through reduced recruitment of *E. superba*.

The fitness consequences of reduced foetus size in *B. m. intermedia* are not known, but compromised feeding by pregnant females is associated with reproductive failure in other marine vertebrates. In Adélie penguins (*Pygoscelis adeliae*), foraging and breeding performance is closely linked to *E. superba* abundance and extent of sea ice (Fraser & Hofmann, 2003). Similarly, Brierly et al. (1997) showed a direct link between fluctuation in *E. superba* abundance and the reproductive performance of Gentoo (*Pygoscelis papua*) and Macaroni penguins (*Eudyptes chrysolophus*). In marine mammals, pregnant females in poor condition limit investment in foetus growth. In the minke whale (*Balaneoptera acutorostrata*) investment in foetal growth by females is in direct proportion to body condition, which is a function of feeding success by pregnant females (Christansen et al., 2014). Near-term foetal body weight was reported to be lower in years with diminished female body condition and feeding conditions in Icelandic populations of *B. physalus* (Lockyer, 1990, 2007). Feeding conditions were also shown to be related to blubber thickness and pregnancy rates in *B. physalus* (Lockyer, 1986; Williams et al., 2013). In southern right whales (*Eubalena australis*) calf abundance between 1997-2013 off the coast of southern Brazil correlated positively with *E. superba* abundance in the waters around South Georgia nine months earlier (Seyboth et al., 2016). Calving success of the same population between 1971-2000 was further shown to be negatively correlated with SST at South Georgia the previous year (Leaper et al., 2006), demonstrating a link between SST, *E. superba* abundance and reproductive success. The impacts of nutritional stress on the development and early survival of offspring of pregnant females are well described in a range of taxa (Lindström, 1999), including marine mammals in which poorly provisioned foetuses experience slow growth and, following parturition, suffer reduced survival rates through an elevated risk of starvation, greater susceptibility to predation and reduced buoyancy (Boltnev et al., 1998; Lockyer, 2007; McMahon et al., 2000, 2003; Reeves et al., 2001).

The data presented here span 30 years, from 1925-1954. In the interval following this period, the extent of Antarctic sea ice and duration of the sea-ice season have declined sharply, in association with elevated SST in the Southern Ocean (de la Mare, 1997; Jacobs & Cosimo, 1997; Smith & Stammerjohn, 2001), with well-characterised negative effects on *E. superba* abundance (Atkinson et al., 2004; Brierly et al., 2002). The impact of these trends on the fitness of *B. m. intermedia*, have hitherto not been demonstrated, though several authors have considered the likely knock-on effects of declining *E. superba* stocks on baleen whale populations (Braithwaite et al., 2015; Brierly et al., 1997, 2002). The implications of the present analysis are that the fitness of *B. m. intermedia* will be compromised by reduced *E. superba* stocks through a negative effect on foetal growth, undermining any positive density-dependent effects through a ‘krill surplus’ at low *B. m. intermedia* abundance. This finding offers a potential explanation for the failure of *B. m. intermedia* to show recovery to anything approaching pre-exploitation levels, with reduced food availability in Antarctic waters predicted to result in under-sized calves with low predicted survival rates. With the prospect of a 95% decline in *E. superba* stocks in regions of the Southern Ocean over the coming century in response to an increase of 1-2 °C in SST (Murphy et al., 2007), strong recovery of *B. m. intermedia* populations, in consequence, appears doubtful.

Because foetuses were not aged, the results presented here could be explained through environmental effects on the time of conception in *B. m. intermedia*, rather than effects on foetal growth rates. Thus, if conception dates were later in years with high SST in the Southern Ocean the same effect would be observed, with smaller foetuses, on average, on a given day of the year, but unrelated to the nutritional state of the female. Model estimates of foetus size-at-age closely matched observed data, though with lower variance (Fig. 3). Some of this difference in variance could be attributed to inaccuracies in the measurement and recording of foetal lengths and with variation in pregnant female condition and foetal growth rates (Frazer & Huggett, 1973). However, an additional explanation for this disparity in predicted and observed size-at-age could be a wider window in which conceptions occurred than assumed. Despite this disparity, there was no evidence for a directional effect of time of conception as a function of environmental conditions in the data (Fig. 4), indicating that model predictions did not differ systematically from observed as a function of temperature. There is also no obvious mechanism by which conception dates might be influenced by SST in the Southern Ocean, since breeding in *B. m. intermedia* takes place at low latitudes during the preceding winter (Roston et al., 2013), though it may influence ovulation. In addition, the data showed clear evidence of density dependence in foetus growth, with a positive relationship between foetus size and cumulative *B. m. intermedia* catch, which indicates foetal sensitivity to female feeding conditions as predicted.

Another potential caveat to this study stems from a temporal shift in whalers taking *B. m. intermedia* to the capture of the smaller pygmy blue whale (*B. m. brevicauda*). However, this switch came in the early 1960s, approximately a decade after the period for which data are presented here (Branch, 2007; Branch et al., 2004). It is notable that pregnant female body size showed no trend, either upward or downward, over the period for which data are presented (Fig. 5), suggesting that pregnant *B. m. brevicauda* did not make a significant contribution to the catch, if at all. Similarly, while the impact of commercial fisheries for finfish and *E. superba* on the trophic structure of the Southern Ocean have been substantial (Hill, Murphy, Reid, Trathan, & Constable, 2006), the onset of commercial fishing in the Southern Ocean came over a decade after the end of the period for which data are presented (Kock, 1994).

In conclusion, the analysis presented here provides circumstantial evidence for a negative relationship between *B. m. intermedia* foetus size and antecedent winter SST in the Southern Ocean. The influence of winter temperature on foetus size is most likely mediated by the positive effect of the extent and duration of winter sea ice on *E. superba* abundance. The effect of winter temperature was shown to interact with *B. m. intermedia* abundance, with a positive density-dependent effect of a ‘krill surplus’ at low whale population sizes lost at elevated winter SST. The effect of predicted SST rises of 1-2 °C in the Southern Ocean (Murphy et al., 2007) are anticipated to compromise the growth of *B. m. intermedia* foetuses, with implications for the capacity of the sub-species to recover from overexploitation.

## Acknowledgements

I am grateful to Andy Brierley, Phil Hammond and Rowena Spence for comments and to John Smith for meticulous data compilation.

## Data Accessibility

Whaling data came from the IWC (www.iwc.int) and are open access. Temperature anomaly data came from the HadISST1 data set compiled by the UK Met Office Hadley Centre (www.metoffice.gov.uk/hadobs/hadisst).

